# Evolutionary thrift: mycobacteria repurpose plasmid diversity during adaptation of type VII secretion systems

**DOI:** 10.1101/067207

**Authors:** Tatum D. Mortimer, Alexandra M. Weber, Caitlin S. Pepperell

## Abstract

Mycobacteria have a distinct secretion system, termed type VII (T7SS), which is encoded by paralogous chromosomal loci (ESX) and associated with pathogenesis, conjugation, and metal homeostasis. Evolution of paralogous gene families is of interest because duplication is an important mechanism by which novel genes evolve, but there are potential conflicts between adaptive forces that stabilize duplications and those that enable evolution of new functions. Our objective was to delineate the adaptive forces underlying diversification of T7SS. Plasmid-borne ESX were described recently, and we found evidence that the initial duplication and divergence of ESX systems occurred on plasmids and was driven by selection for advantageous mutations. Plasmid conjugation has been linked to T7SS and type IV secretion systems (T4SS) in mycobacteria, and we discovered that T7SS and T4SS genes evolved in concert on the plasmids. We hypothesize that differentiation of plasmid ESX helps to prevent conjugation among cells harboring incompatible plasmids. Plasmid ESX appear to have been repurposed following migration to the chromosome, and there is evidence of positive selection driving further differentiation of chromosomal ESX. We hypothesize that ESX loci were initially stabilized on the chromosome by mediating their own transfer. These results emphasize the diverse adaptive paths underlying evolution of novelty, which in this case involved plasmid duplications, selection for advantageous mutations in the mobile and core genomes, migration of the loci between plasmids and chromosomes, and lateral transfer among chromosomes. We discuss further implications for the choice of model organism to study ESX functions in *Mycobacterium tuberculosis*.

## Introduction

Gene duplications are an important mechanism by which novel gene functions evolve (Zhang 2003). Duplications have been shown to occur frequently during experimental evolution of bacterial populations and can be adaptive, for example in producing antibiotic resistance (Sandegren & Andersson 2009). However, most duplications are transient, due to their intrinsic instability and associated fitness costs, as well as general mutational biases toward deletion (Sandegren & Andersson 2009; Adler et al. 2014).

These observations have led researchers to investigate the selective forces allowing duplicate genes to persist and to diverge from the parent gene (Bershtein & Tawfik 2008; Bergthorsson et al. 2007; Näsvall et al. 2012). ‘Ohno’s dilemma' refers to the potential conflict between selection that stabilizes duplicated genes and that which enables evolution of novel functions (Ohno 1970). Selection that stabilizes the initial duplication is likely to preserve the gene's original function and prevent differentiation from the parent gene, whereas selection for a new function would drive differentiation of the gene copies. Several solutions have been proposed (Bergthorsson et al. 2007; Hittinger & Carroll 2007; Elde et al. 2012) to the problem of how duplicated genes are maintained and allowed to differentiate such that novel functions can evolve.

Bacterial species within the genus *Mycobacterium* have a distinct secretion system, termed the type VII secretion system (T7SS), which is encoded by six paralogous chromosomal loci referred to as ESX (eSX-1, -2, -3, -4, -5, and -4-bis/-4_EvoL_). The ESX loci share a core consisting of 6 genes *(eccB, eccC, eccD, mycP, esxA, esxB);* the loci typically encode an additional 4 genes (a PE, PPE (Bottai & Brosch 2009), *eccA*, and *eccE*) as well as a variable complement of locus specific gene content (Figure 1).

**Fig. 1.**
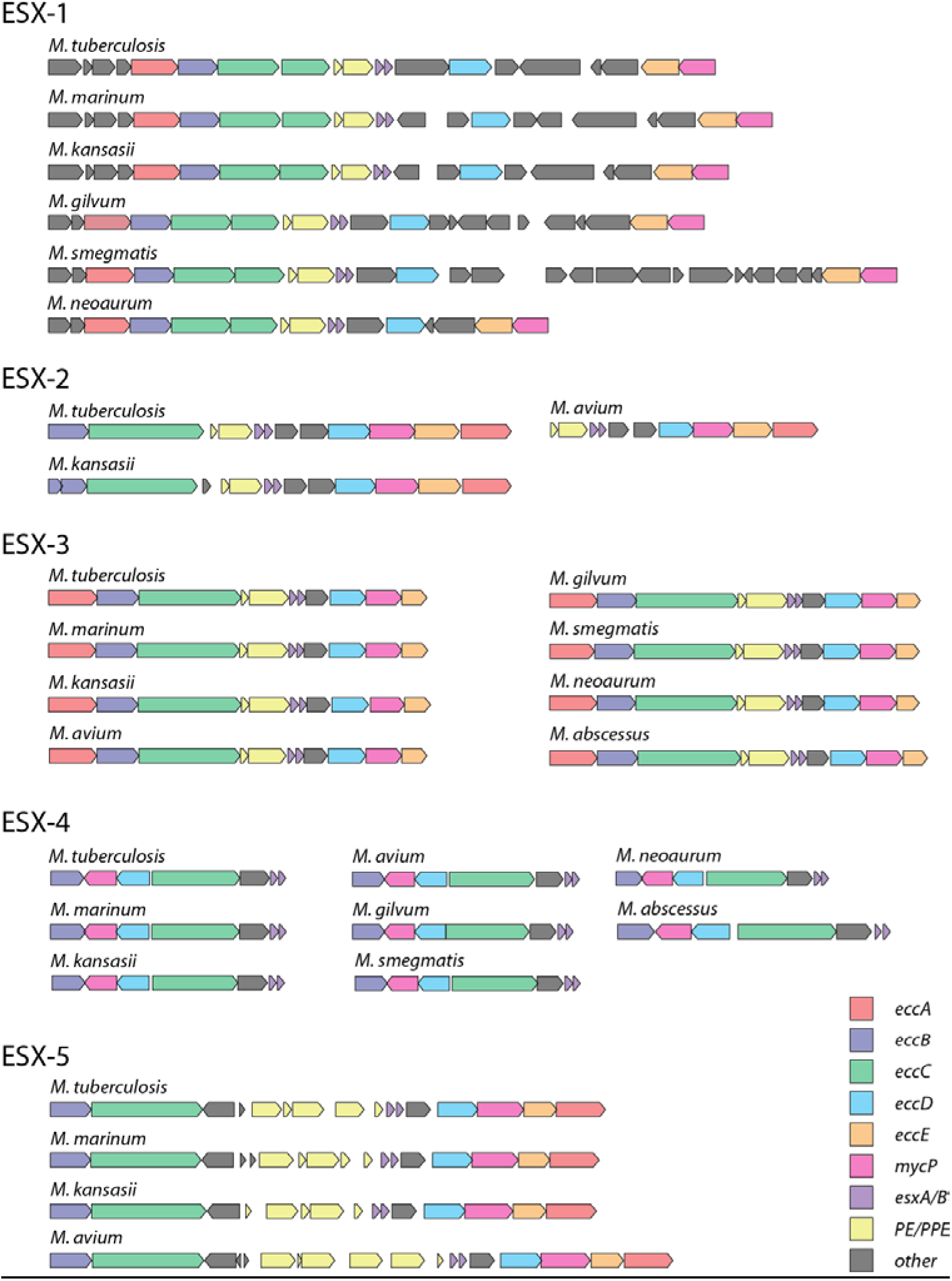
Mycobacterial chromosomal ESX loci. Core gene content in the ESX loci are colored as follows: *eccA-* red, *eccB-* dark blue, *eccC-* green, *eccD-* light blue, *eccE-* orange, *mycP-* pink, *esxA/B-* purple, *PE/PPE-* yellow. Other variable genes in the loci are black. Orthologs and paralogs are based on OrthoMCL (Li et al. 2003b) output. Locus diagrams were made using GenomeTools (Gremme et al. 2013). Each locus has a distinct structure, which developed during adaptation on mycobacterial plasmids and chromosomes (see text for details).

Functions have not been identified for all ESX loci, but the available data indicate that duplicated T7SS loci are associated with diverse functions. ESX-1 is associated with several aspects of virulence in *Mycobacterium tuberculosis,* including growth in macrophages (Stanley et al. 2003; McLaughlin et al. 2007), cytosolic translocation (Houben et al. 2012), and antigen presentation (Sreejit et al. 2014). In *Mycobacterium smegmatis,* a non-pathogenic, environmental mycobacterium, ESX-1 and ESX-4 are involved in distributive conjugal transfer, a mechanism of lateral gene transfer (Flint et al. 2004; Coros et al. 2008; Gray et al. 2013, 2016). ESX-3 is essential for *M. tuberculosis* growth *in vitro* (Sassetti et al. 2003) and is involved in iron acquisition in mycobacteria (Serafini et al. 2009; Siegrist et al. 2009; Serafini et al. 2013). ESX-3 is also thought to contribute to *M. tuberculosis* virulence independent of its role in metal homeostasis (Mehra et al. 2013; Tufariello et al. 2016). ESX-5 has been shown to secrete PE/PPE proteins in *Mycobacterium marinum* (Abdallah et al. 2009) and *M. tuberculosis* (Bottai et al. 2012). The emergence of ESX-5 coincides with the expansion of PE/PPEs in mycobacteria (Pittius et al. 2006). Both ESX-1 and ESX-5 additionally play roles in membrane integrity (Garces et al. 2010; Ates et al. 2015). The function of ESX-2 in mycobacteria is unknown.

The goal of the present study was to delineate the adaptive processes underlying divergence of mycobacterial T7SS, and to define groups of T7SS that are likely to be functionally related. In our analyses of genomic data from 33 mycobacterial species and related Actinobacteria, we found evidence pointing to complex dynamics between the core and mobile genomes underlying adaptation of the paralogous chromosomal ESX loci. Positive selection appears to have played a role in the duplication and divergence of these loci, and we hypothesize about how such selection might operate on plasmids and the chromosome. Loci within groups that diverged from each other because of positive selection are likely to be functionally related and based on our results, we propose model organisms for the study of ESX functions in *M. tuberculosis*.

## Materials and Methods

### Data set

We obtained finished genomes from all available *Mycobacterium* species (n = 30, as of December 2015) and 23 representative Actinobacteria genomes from the National Center for Biotechnology Information (NCBI) database. Accession numbers for these genomes can be found in Table S1. Members of the *M. tuberculosis* complex (MTBC) without finished genomes *(M. caprae, M. pinnipedii, M. orygis)* were assembled by the reference guided assembly pipeline available at https://github.com/tracysmith/RGAPepPipe using *M. tuberculosis* H37Rv as the reference. Briefly, reads were trimmed for quality and adapters using Trim Galore! v 0.4.0 (Kreuger 2013); trimmed reads were mapped to the reference genome using BWA-MEM v 0.7.12 (Li 2013); Picard-tools v 1.138 (https://broadinstitute.github.io/picard/) marked duplicates and added read group information; and variants were called using GATK v 3.4.46 (DePristo et al. 2011).

### Ortholog detection

Genomes were annotated using Prokka v 1.11 (Seemann 2014). We used OrthoMCL v 2.0.9 (Li et al. 2003a) to cluster proteins from these genomes into orthologous groups. Genes known to be located in the ESX loci of *M. tuberculosis* H37Rv were obtained from (Bitter et al. 2009). Orthologous groups containing any of the genes in ESX loci of *M. tuberculosis* were identified. ESX loci were identified as at least three orthologs of genes present in *M. tuberculosis* ESX loci, in close proximity to one another in the genome. Identification of ESX loci was confirmed by phylogenetic analysis of conserved genes as described below.

### ESX loci and core genome alignment

Protein sequences from paralogs and orthologs of genes present in the majority of ESX loci in mycobacteria (eccA, *eccB, eccC, eccD, eccE, mycP)* were aligned with MaFFT v 7.245 (Katoh & Standley 2014), low quality alignment columns were identified and removed using GUIDANCE v 2.01 (Sela et al. 2015), and trimmed alignments were concatenated to produce an alignment of ESX loci. We additionally identified orthologous groups present in every genome only one time as the core genome. Alignments of core proteins produced with MAFFT were concatenated for phylogenetic analysis. Scripts used to automate OrthoMCL analysis and alignment can be found here: https://github.com/tatumdmortimer/core-genome-alignment.

### Plasmid assembly and annotation

Since there are few finished, mycobacterial plasmid sequences available that contain ESX loci, we screened publicly available sequence data for evidence of plasmid-borne ESX. Sequence reads identified as *Mycobacterium,* excluding those belonging to the MTBC or *Mycobacterium leprae,* which are not known to harbor plasmids, were downloaded and assembled using plasmidSPAdes v 3.5.0 (Antipov et al. 2016). Resulting plasmid contigs were annotated using Prokka v 1.11 (Seemann 2014).

Plasmids with at least one annotated ESX gene were chosen for further quality control processing, including checking for at least three ESX genes, checking that all ESX genes were on the same component when multiple components were assembled, and ruling out chromosomal ESX loci misidentified as plasmid-borne. In total, we downloaded and assembled reads from 1300 *Mycobacterium* strains, resulting in 732 strains with assembled plasmids. We sampled at least one strain from 67% of named *Mycobacterium* species with sequence data available in NCBI, and 50% of *Mycobacterium* strains without a species designation. The majority of nontuberculous mycobacteria reads available in NCBI are *M. abscessus (n =* 1990), and we assembled 20% of these strains. Two hundred and forty eight strains contained a plasmid with at least one ESX gene, and 16 plasmids passed all quality control checks (Table S2). Final identification and alignment of ESX loci in these assembled plasmids as well as publicly available plasmid sequences (Table S3) was performed as described above for the chromosomal loci. While *M. ulcerans* plasmids were not included in the downstream analyses because they did not contain a complete ESX locus, we did create a core gene alignment (21 genes) and phylogeny in a sample (n = 7) of the total *M. ulcerans* plasmids assembled.

### Phylogenetic analysis

We performed all phylogenetic analyses using RAxML v. 8.2.3 (Stamatakis 2014). The best protein model was determined automatically using the -m PROTGAMMAAUTO option. The best-scoring maximum likelihood tree was calculated from 20 trees, and bootstrap values were calculated using the autoMR bootstrap convergence criteria. We used Dendroscope v 3 (Huson & Scornavacca 2012) and ggtree (Yu et al. 2016) for tree visualization and editing. Phylogenetic networks were created using Splitstree 4 (Huson & Bryant 2006), and we used the PHI test (Bruen et al. 2006) to assess the presence of recombination in the alignments. In order to address the congruence of core plasmid genes, we performed Bayesian phylogenetic analysis using MrBayes v 3.2.5 (Ronquist & Huelsenbeck 2003) and visualized tree clusters using Treescape (Kendall & Colijn 2016). MrBayes analysis was run for 1,000,000 generations for each gene, and trees were sampled every 500 generations. We discarded the first 25% of trees as burn in, randomly sampled 200 trees from each gene, and performed pairwise calculations of the Kendell Colijn metric and multidimensional scaling in Treescape. This analysis was performed on a subset of plasmids encoding the gene *nrdH,* as there were no genes common to all plasmids outside of T7SS and T4SS.

### Selection analysis

We used the aBSREL method implemented in HyPhy (Smith et al. 2015) to test for episodic directional selection in a tree of mycobacterial ESX loci. The method initially assumes that each branch in the phylogeny can be modeled with only one rate class, and rate classes are added to each branch in a step-wise manner only if there is an improvement in the likelihood of the data given the model. The resulting model allows rate variation across branches and sites. Additionally, this method identifies branches on the phylogeny where there is evidence for a proportion of sites to be modeled with an ω (dN/dS) greater than 1 (indicative of positive selection). We tested all branches for positive selection, and branches with a p-value < 0.05 after the Holm-Bonferroni multiple testing correction were considered to have statistically significant evidence for directional selection. Nucleotide sequences from ESX genes were aligned with MAFFT, trimmed with Guidance, and concatenated for input into the HyPhy analysis. Additionally, a nucleotide alignment was created using translatorX (Abascal et al. 2010), which back-translates an amino acid alignment to preserve the reading frame of codons, and trimmed with Gblocks v 0.91b (Castresana 2000). Both alignments were used for maximum likelihood phylogenetic inference with RAxML and HyPhy analysis.

### Data availability

Unless stated otherwise above, all scripts and data, including text files for supplementary tables, used in these analyses are available at https://github.com/tatumdmortimer/t7ss.

## Results

### Rapid expansion of plasmid T7SS followed by migration to the chromosome

A map of the chromosomal ESX loci is shown in Figure 1. Figure 2 shows a core genome phylogeny of 56 species of Actinobacteria along with a presence/absence matrix of associated chromosomal T7SS. Our analyses are consistent with an initial emergence of the FtsK/WXG100 gene cluster on the chromosome (as proposed in (Pallen 2002)), followed by ESX-4-bis and ESX-4, with subsequent duplications giving rise to ESX-3, ESX-1, ESX-2, and ESX-5 (as proposed in (Pittius et al. 2006; Dumas et al. 2016; Newton-Foot et al. 2016)). Interestingly, the loci have been lost from the chromosome on several occasions. For example, ESX-2 was lost in the common ancestor of *M. marinum, M. liflandii,* and *M. ulcerans,* and ESX-1 has been lost in *M. sinense, M. avium* and related species, as well as from *M. ulcerans*.

**Fig. 2.**
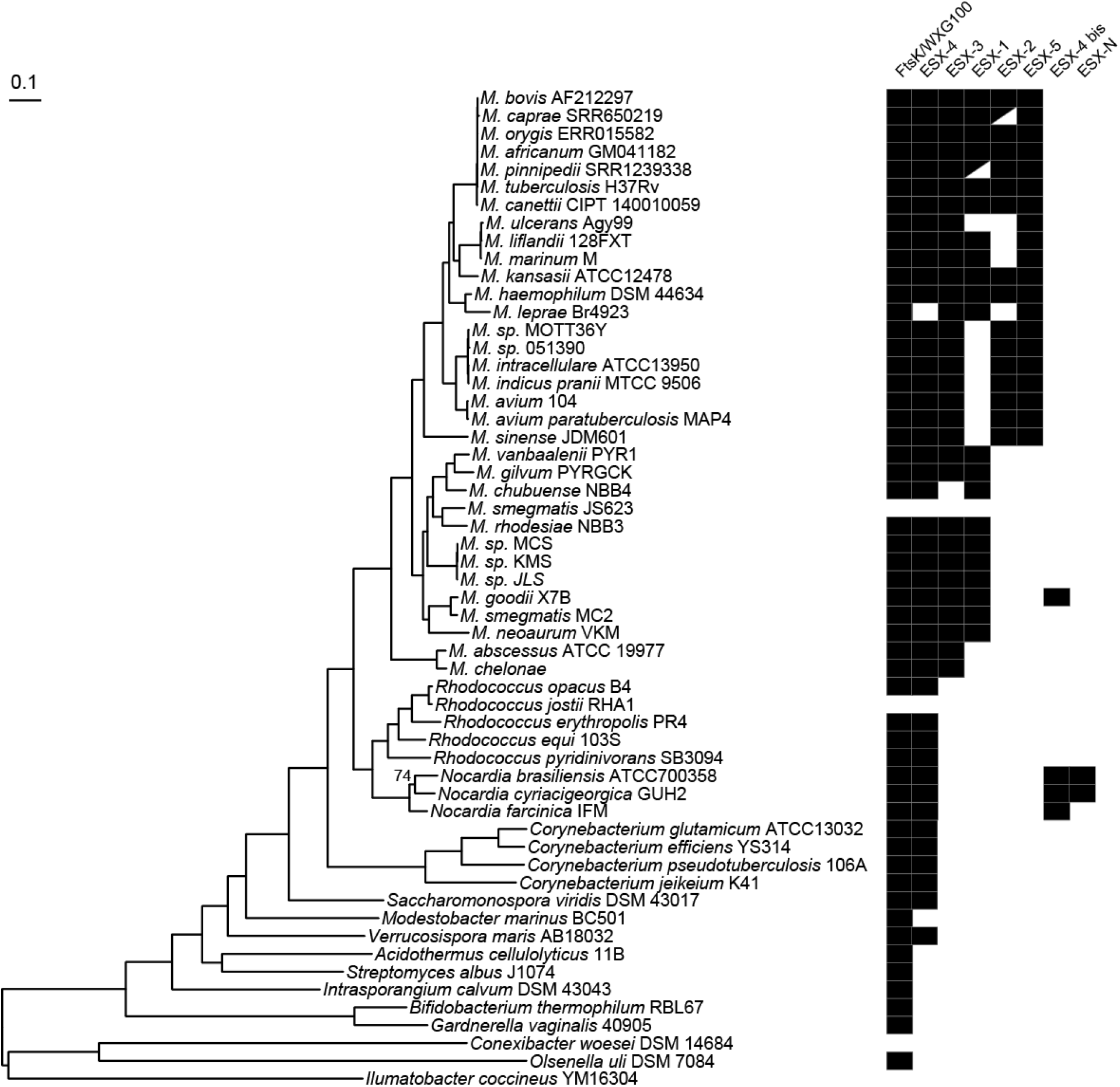
Maximum likelihood phylogeny of Actinobacteria with presence/absence matrix of type VII secretion system loci. RAxML was used for phylogenetic inference of the Actinobacteria core genome alignment (concatenated amino acid alignments of genes (n = 171) present in all genomes without duplications). The phylogeny is midpoint rooted, and branches without labels have a boostrap value of 100. Presence of ESX loci is indicated with black boxes. We have abbreviated the genus *Mycobacterium* in the tip labels. Some *M. tuberculosis* complex (MTBC) species have characteristic deletions located in ESX loci. Partially deleted ESX loci are represented by black triangles. *M. caprae* has a deletion in ESX-2 spanning PE/PPE, *esxC, espG2, Rv3888c, eccD2,* and *mycP2. M. pinnipedii* has a deletion in ESX-1 spanning PE/PPE and a portion of *eccC1b*. Patterns of ESX presence/absence are consistent with an initial emergence of the FtsK/WXG100 gene cluster, followed by ESX-4 bis and ESX-4, with subsequent duplications giving rise to ESX-3, ESX-1, ESX-2 and ESX-5.

Figure 3 shows a network of plasmid and chromosomal ESX loci (see also Figure S1). The network has a pronounced star-like configuration, consistent with rapid diversification of these loci. This pattern is particularly evident when the plasmid loci are considered separately (Figure 4).

**Fig. 3.**
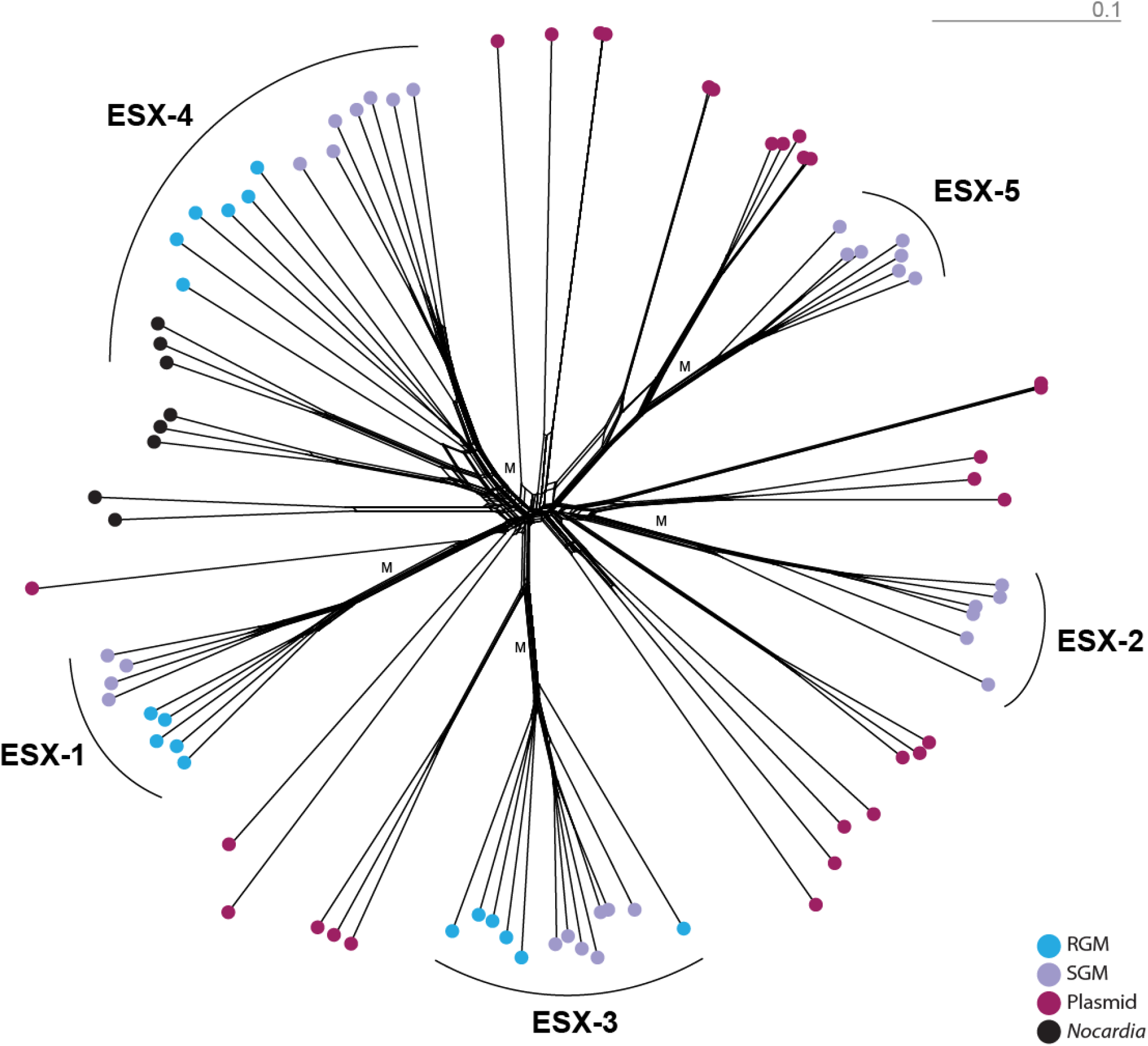
Network of ESX loci in mycobacteria, *Nocardia,* and mycobacterial plasmids. The network was created in SplitsTree4 from a concatenated alignment of *eccA, eccB, eccC, eccD, eccE,* and *mycP*. Light blue dots correspond to ESX loci from rapid growing mycobacterial chromosomes (RGM), light purple dots correspond to ESX loci from slow growing mycobacterial chromosomes (SGM), magenta dots correspond to ESX loci from mycobacterial plasmids, and black dots correspond to ESX loci from *Nocardia* chromosomes. The earliest branching lineages are all plasmid-associated, suggesting that the ancestral ESX locus was plasmid-borne (putative location of migration events to the chromosome marked 'M' on the network). The PHI test was insignificant (p = 1.0) for this alignment, indicating that there was no evidence for intra-locus recombination. A version of this figure with tip labels is available in the supplement (Figure S1).

**Fig. 4.**
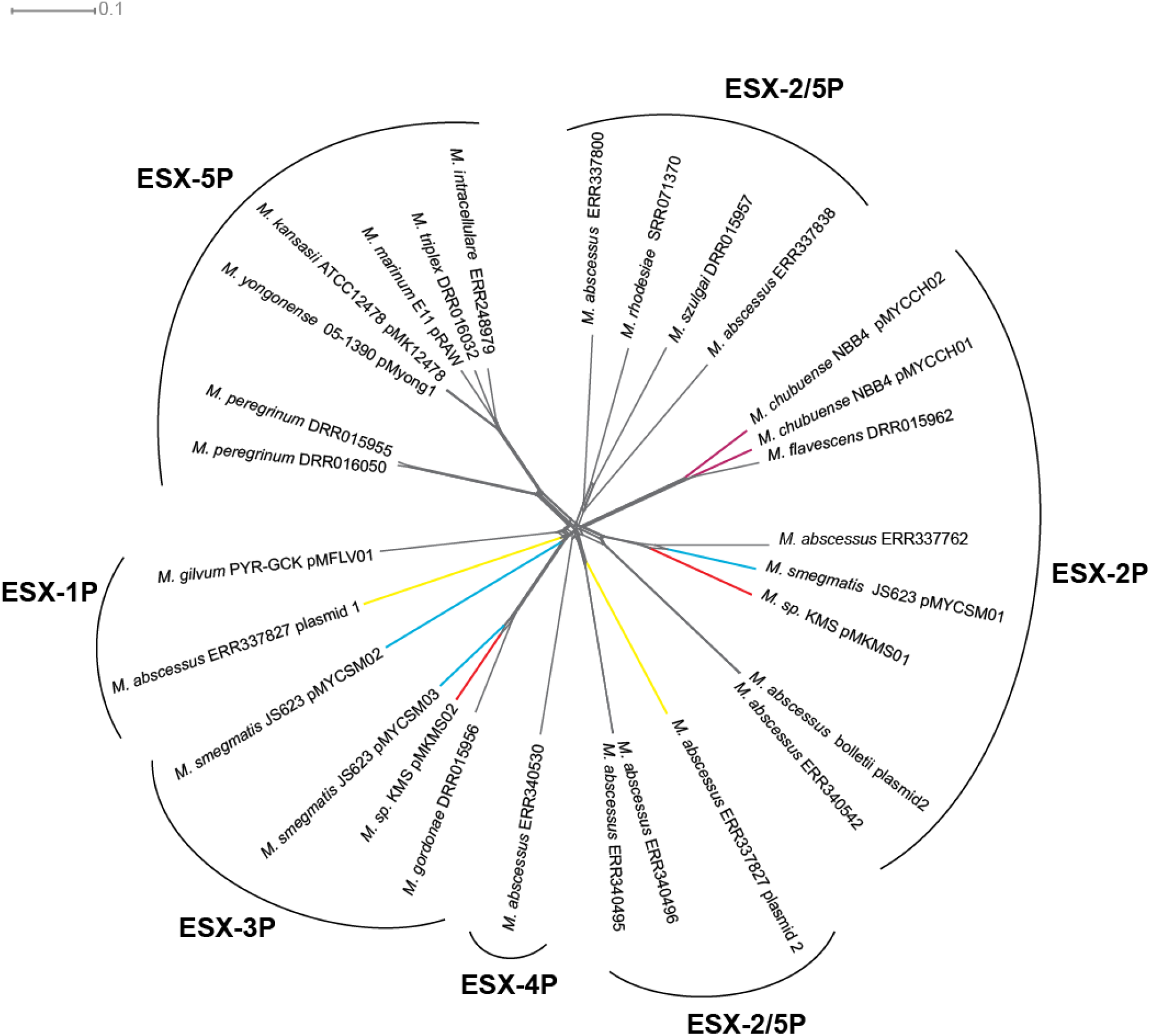
Network of plasmid-borne ESX loci. The network was created in SplitsTree4 from a concatenated alignment of *eccA, eccB, eccC, eccD, eccE,* and *mycP* from ESX loci encoded on mycobacterial plasmids. The star-like appearance of the network is consistent with rapid diversification of this gene family on the plasmids. Some bacterial strains harbored multiple plasmids, and these are indicated with colored branches. The phylogenetic relationships among plasmid ESX loci do not follow the core genome phylogeny, and plasmids with divergent ESX loci can be found within the same host species or even the same cell. This suggests that plasmid ESX diversification has not been shaped by adaptation to bacterial host species. We did not find evidence of intra-locus recombination in this alignment with the PHI test (p = 1.0).

Plasmid ESX that are basal to chromosomal ESX-1, -3, -2 and -5 have been described previously (Dumas et al. 2016; Newton-Foot et al. 2016). We identified several new, plasmid-borne ESX lineages, including a plasmid lineage that is basal to ESX-4 (Figures 3 and 5B). The most parsimonious explanation of these observations is that the common ancestor of ESX-1 through -5 was plasmid-associated, that duplication of the ESX loci occurred on plasmids, and that extant chromosomal loci all result from transfers from plasmid to chromosome (Figure 5A). It is possible that the common ancestor of ESX 1-5 was chromosomal, as suggested by Newton-Foot and Dumas et al; however, this scenario would require more migration events.

**Fig. 5.**
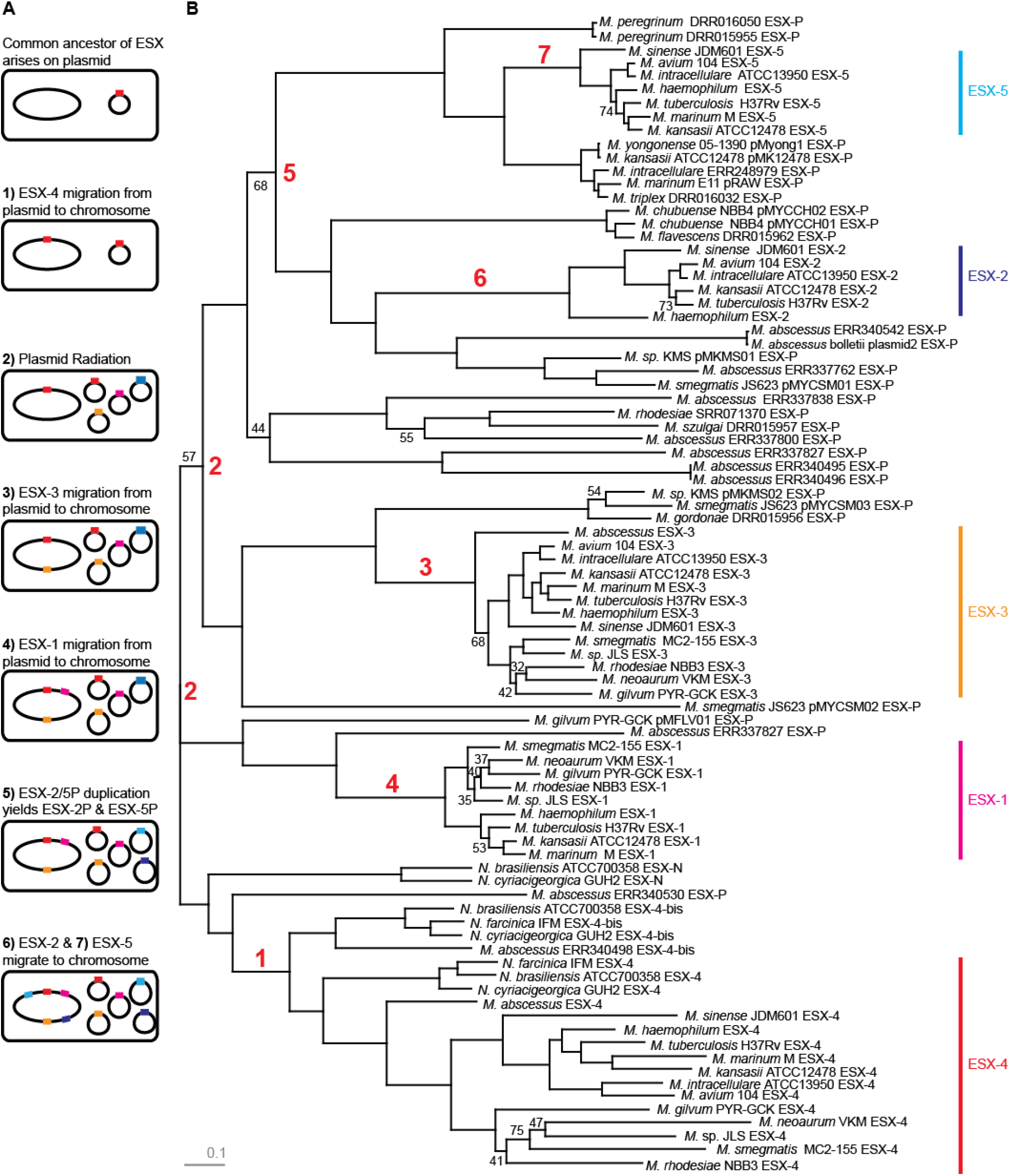
ESX plasmid-mediated duplication and migration to the chromosome. A) Simplified schematic of major steps in the evolutionary history of mycobacterial ESX loci. ESX loci are colored as follows: Ancestral/ESX-4: red, ESX-3: orange, ESX-1: pink, ESX-2: light blue, ESX-5: dark blue. B) Maximum likelihood phylogeny of ESX loci *(eccA, eccB, eccC, eccD, eccE,* and *mycP)* in mycobacteria, *Nocardia,* and mycobacterial plasmids. Branches without black labels have a bootstrap value greater than 75. Red labels correspond to events presented in the schematic. ESX-N, found on the chromosomes of some *Nocardia* species, appear to have been recently transferred from a plasmid (see text). ESX-N and other plasmid-associated ESX are basal to chromosomal ESX 1-5. This suggests that their common ancestor was plasmid-borne and that extant chromosomal loci trace to migrations from plasmid to chromosome. The model outlined here is highly simplified: for example, there were likely several migrations of ESX-4 like loci to the chromosome (step 1 in the schematic) and the chromosomal loci show a mixture of vertical and horizontal inheritance (details in text).

Chromosomal ESX-5, found exclusively among slow growing mycobacteria (SGM), is related to plasmid loci from both SGM and rapid growing mycobacteria (RGM), suggesting that ESX-5 like loci diversified on RGM and SGM-associated plasmids prior to their migration to the chromosome of SGM. Although we did not identify any complete ESX-2 like loci in SGM plasmids, the incomplete ESX locus we identified in *M. ulcerans* (SGM) is most closely related to ESX-2, suggesting that the same may be true of ESX-2.

The two most basal mycobacterial species, *M. abscessus* and *M. chelonae,* have a chromosomal ESX-3 locus, but not an ESX-1 locus. ESX-1 is, however, basal to ESX-3 on the ESX phylogeny, on a branch with low bootstrap values (57%, Figure 5B). We speculate that this conflict - i.e. between the species ranges and phylogenetic positions of ESX-1 and ESX-3 - as well as the uncertainty in the phylogeny is due to the plasmid-borne ancestor of ESX-1 having emerged earlier than ESX-3, but ESX-3 being first to migrate to the chromosome.

### Lateral transfer of T7SS: migrations between plasmids and the chromosome, transfer among chromosomes, intra-locus recombination

A phylogeny of ESX-4 and related loci is shown in Figure 6. ESX-N, found on the chromosome of *Nocardia brasiliensis* and *N. cyriacigeorgica,* pairs with a plasmid-associated ESX locus, and is basal to ESX-4 and related ESX from a range of actinobacterial species. ESX-4 and ESX-4-bis appear to be fixed among Nocardia species and are stably associated with flanking gene content, suggesting vertical inheritance in the genus. ESX-N, by contrast, is variably present among sampled Nocardia species, and we found it to be associated with variable flanking gene content (Figure S2). We also found ESX-N in association with T4SS genes and other gene content otherwise specific to plasmids. We hypothesize that the ESX-N loci were horizontally transferred from an unsampled (or extinct) plasmid.

**Fig. 6.**
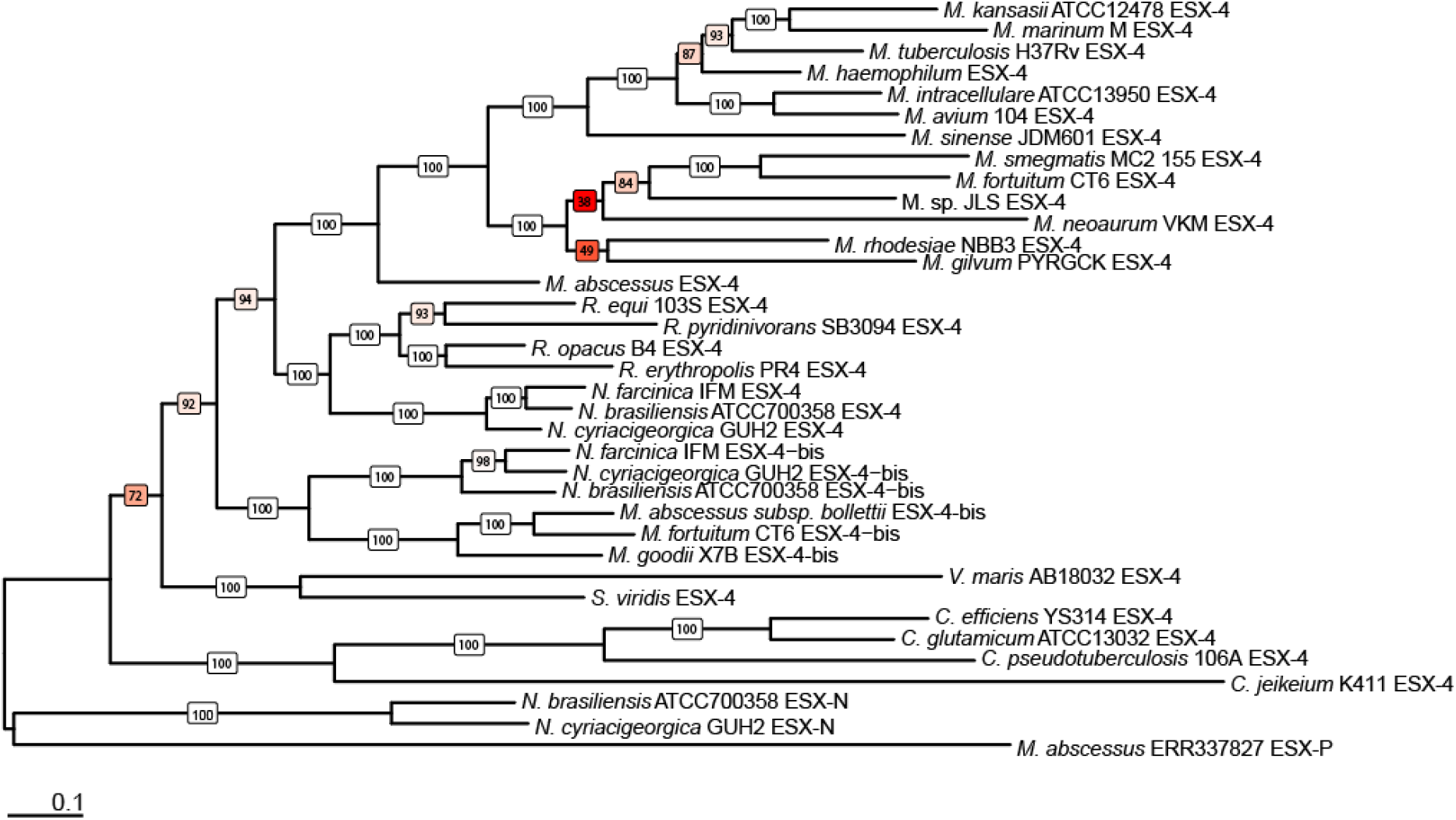
Maximum likelihood phylogeny of ESX-4 in Actinobacteria. The phylogeny is rooted using ESX-N and a basal plasmid-borne ESX locus. Bootstrap values are colored based on support (white = 100, red = lowest support). The location of chromosomal *Corynebacterium* ESX-4 and *M. goodii* ESX-4-bis are in conflict with the core genome phylogeny (Figure 2). In the core genome phylogeny, *Corynebacterium* is more closely related to *Nocardia* and *Rhodococcus* than *Verrucosispora* or *Saccharamonaspora*. However, in the ESX-4 phylogeny, this relationship is reversed. Based on the relationship of *Mycobacterium* species in the core genome phylogeny, we would expect *M. goodii* ESX-4-bis and *M. fortuitum* ESX-4-bis to be more closely related to one another than either is to *M. abscessus* (the most basal *Mycobacterium* species). These conflicts suggest that chromosomal ESX-4 like loci have been laterally transferred among species. Genus names have been abbreviated, but full length names can be found in the core genome phylogeny (Figure 2).

The chromosomal ESX-4 phylogeny is not concordant with that of the core genome (e.g., the placement of corynebacteria), which suggests that the locus was laterally transferred during divergence of the Actinobacteria. The patchy distribution of ESX-4-bis among mycobacterial species, as well as branching patterns among these loci, suggest ESX-4-bis has also been laterally transferred (between chromosomes and/or migrated between plasmids and chromosomes) on a few occasions in the genus. The ESX-4-bis locus in *M. goodii* includes *espI,* which is not found in other chromosomal ESX-4 loci but is part of the plasmid core genome (discussed further below). This suggests the locus, like ESX-N, was transferred relatively recently from a plasmid.

Although broad groupings seen on the core genome (e.g., separation of slow-growing from rapid-growing mycobacteria) are reflected in the phylogeny of the combined ESX loci (Figure 5B), the branching within these groups does not always reflect the patterns of the core genome. Branching patterns within these groups were sensitive to the sampling scheme and alignment, whereas internal branching patterns were stably supported across multiple analyses. This pattern could be due to a lack of fine scale phylogenetic signal in the gene content shared among ESX loci, or to lateral transfer of the loci. To help distinguish between these possibilities, we created an alignment and phylogeny of only ESX-5, which contains information from two additional genes. We found that the phylogenetic uncertainty remained (Figure S3). This suggests that T7SS were laterally transferred among mycobacterial species during their divergence, contributing to both phylogenetic uncertainty and conflicts with the core genome phylogeny.

There are few reticulations in the ESX networks (Figures 3 and 4), suggesting that within-locus recombination has not played a major role in adaptation of these loci. The PHI test for recombination (Bruen et al. 2006) was not significant (p=1.0) for an alignment of chromosomal and plasmid-associated loci, nor for the plasmid-associated loci considered separately. The PHI test was, however, significant (p=1.2 × 10^-5^) for the ESX-5 alignment, suggesting that within-locus recombination has occurred among more closely related loci.

### What drove differentiation of T7SS?

Diversification of T7SS loci could have been driven by neutral or selective forces: the solutions proposed to Ohno's dilemma have incorporated both neutral and Darwinian evolution following gene duplication (Zhang 2003). We tested for episodic directional (positive) selection in the eSx phylogeny using HyPhy (Figure 7). Branches under selection in this model mark periods during which there is evidence of advantageous mutations driving divergence from an ancestral state. We found evidence of positive selection at each ESX duplication event (internal branches connecting duplicate loci), and the branches leading to chromosomal ESX loci in all cases showed evidence of positive selection. Since migration events could have occurred anywhere along the branches connecting plasmid-associated and chromosome-associated nodes, selection associated with this transition may have acted on plasmid ESX, chromosomal ESX or both. The proportion of sites under positive selection varied substantially, with the highest proportion associated with long, plasmid associated tips. These results were replicated across multiple analyses, including different sampling schemes and alignment trimming methods (Figures S4 and S5).

**Fig. 7.**
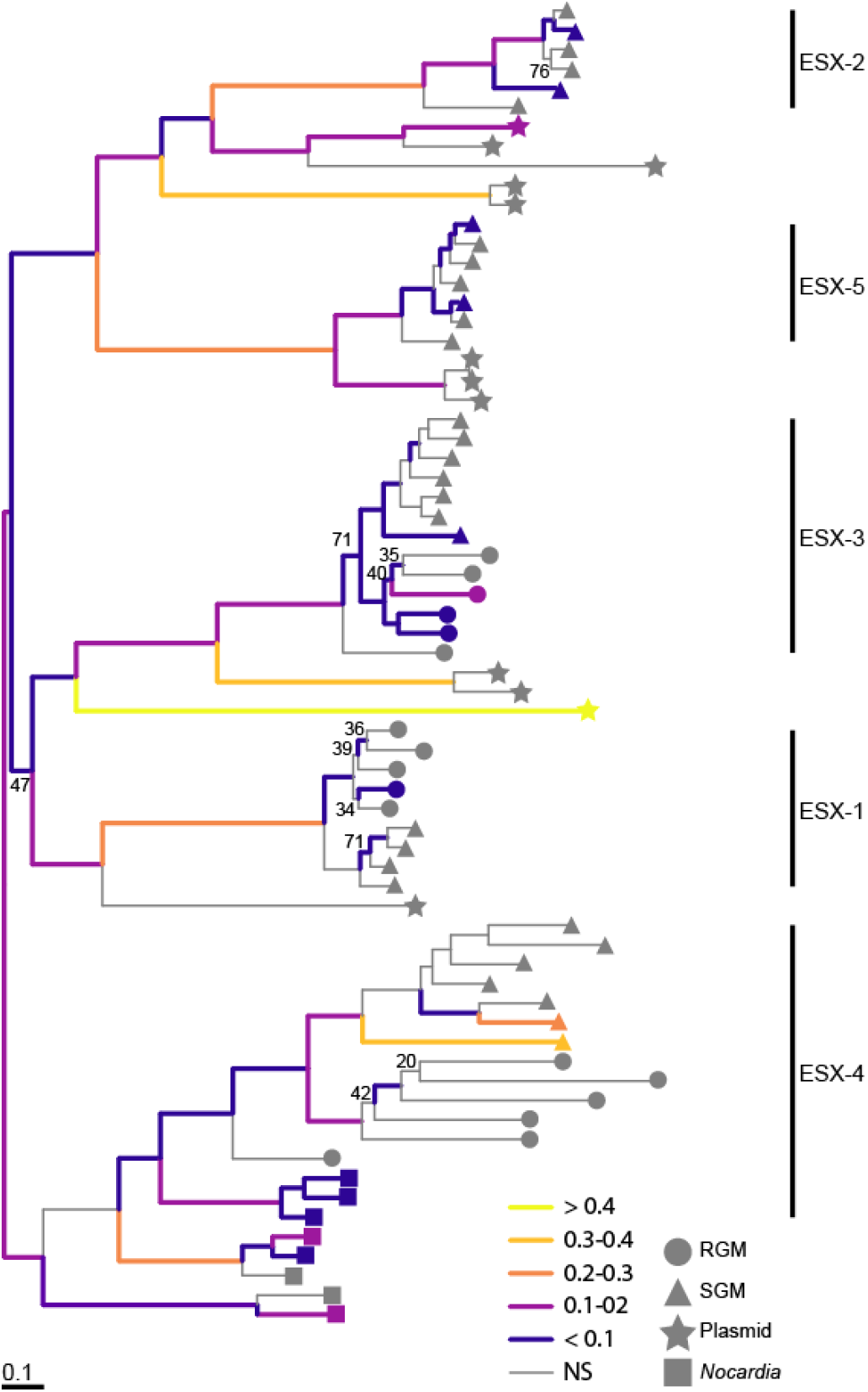
Episodic directional selection during the evolution of ESX loci. Maximum likelihood phylogeny inferred using RAxML from a concatenated alignment of *eccA, eccB, eccC, eccD, eccE,* and *mycP*. In order to minimize potential effects of misalignment on inference of selection, only data from finished genomes were included in this analysis: see Figures 3 and 5B for network and phylogenetic analyses of the complete dataset. Plasmid associated taxa for ESX-4, ESX-2 and ESX-5 are not shown on this phylogeny for this reason. Branches in this phylogeny without labels have a bootstrap value greater than 75. We used the aBSREL test implemented in HyPhy to identify branches with significant evidence (p < 0.05) of episodic directional selection; these branches are colored based on the proportion of sites affected by positive selection (ω > 1). Circles correspond to eSx loci from rapid growing mycobacterial chromosomes, triangles correspond to ESX loci from slow growing mycobacterial chromosomes, stars correspond to ESX loci from mycobacterial plasmids, and squares correspond to ESX loci from *Nocardia* chromosomes. A version of this figure is included in the supplement (fig S4) that shows the tips labeled with species names. There is evidence of directional (positive) selection at each duplication event (short internal branches), as well as on the branches leading to the extant chromosomal loci.

**Fig. 8.**
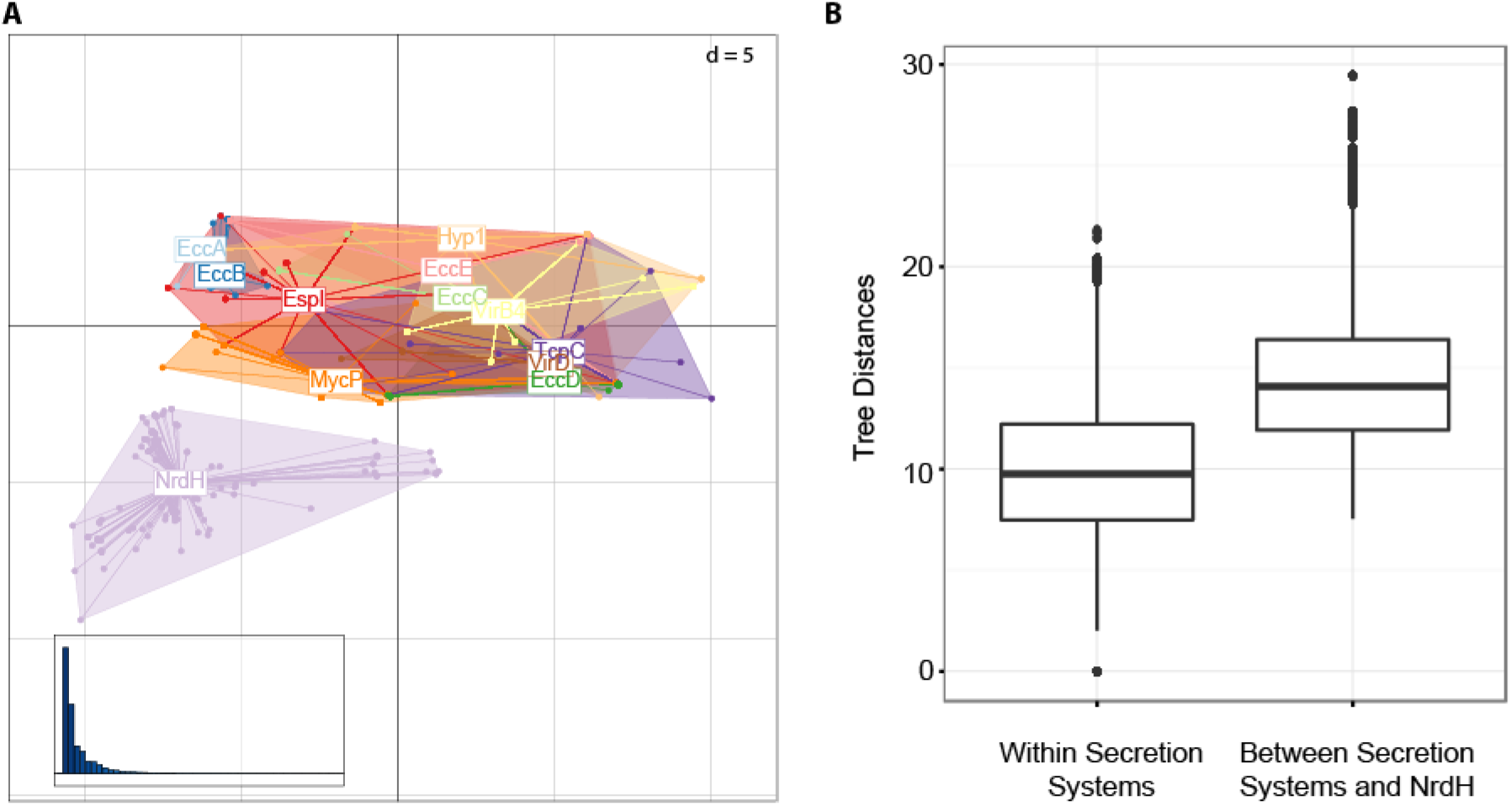
Congruence of tree topologies of plasmid encoded secretion systems. Bayesian phylogenetic analysis was performed in MrBayes using amino acid alignments of EccA, EccB, EccC, EccD, EccE, MycP, VirB4, VirD, TcpC, EspI, NrdH, and a hypothetical protein (Hyp1) encoded proximal to known T4SS genes. A) We used TreeScape to calculate the Kendell Colijn metric between pairs of trees and perform multidimensional scaling (MDS). Clusters of trees are visualized as a scatterplot of the first and third principal components from the MDS. The inset bar chart is a scree plot showing the eigenvalues for the principal components. The T4SS and T7SS gene trees overlap in the MDS, whereas topologies of NrdH gene trees are incongruent with those of T4SS and T7SS. This suggests that plasmid-encoded T4SS and T7SS have co-diverged during their evolutionary history and that they have evolved independently of other gene content on the plasmids. B) Kendell Colijn distances among secretion system gene trees and between secretion system and NrdH gene trees. The means of these distributions are significantly different (p < 2.2 × 10^-16^) according to a Mann-Whitney U test.

**Fig. 9.**
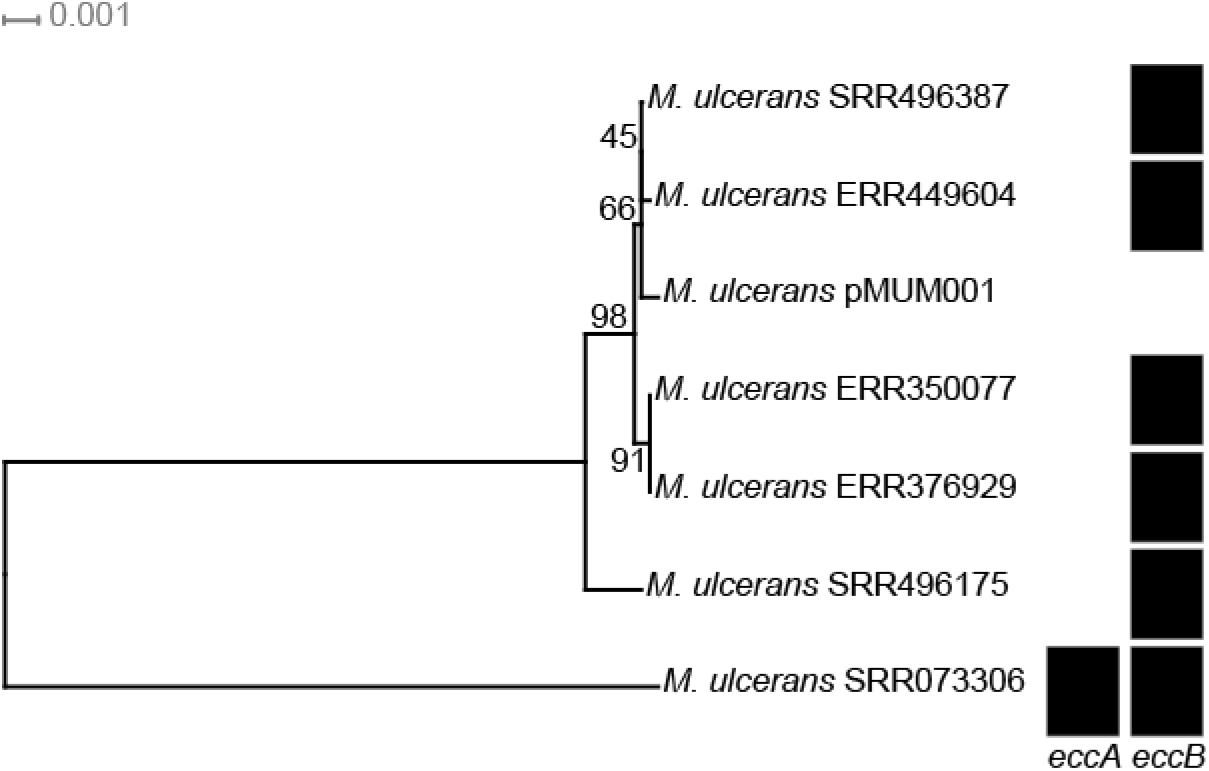
Core gene phylogeny of *Mycobacterium ulcerans* plasmids and presence/absence of T7SS genes. RAxML was used for phylogenetic inference from a core gene alignment (concatenated amino acid alignments of genes (n = 21) present in all *Mycobacterium ulcerans* plasmids without duplications). The phylogeny is midpoint rooted. Presence of ESX genes is indicated with black boxes. ESX genes found on *M. ulcerans* plasmids are most closely related to ESX-2. The most basal *M. ulcerans* plasmid encodes both *eccA* and *eccB*. Most *M. ulcerans* plasmids only encode *eccB,* and some have lost all ESX genes (e.g. pMUM001). The locus shows other signs of degradation (discussed in the text). We hypothesize that selection to maintain the conjugation machinery of these plasmids has been relaxed as a result of host selection for its other gene content, likely mycolactone.

Summarizing the results outlined above, the ESX gene family expansion likely occurred on plasmids, and this diversification appears to have been driven by selection for advantageous mutations. Given that divergence of T7SS loci appears to have been driven by selection for advantageous mutations, we were curious about how such selection might operate. A simple explanation for the segregation of diverse plasmid ESX lineages would be that the plasmids diverged in response to divergence of their host mycobacterial species. In this case, we would expect to observe congruence between the plasmid ESX phylogeny and the host genome phylogeny. However, in this sample of plasmids harboring ESX, the phylogenetic signals are clearly at odds with those of the host genomes (Figure 4): for example, *M. kansasii* pMK12478 pairs with *M. yongonense* pMyong1, rather than *M. marinum* pRAW. There are also multiple divergent plasmid ESX lineages associated with the same host species (e.g., *M. abscessus)* or the same host cell (Figure 4).

Another possible explanation of the plasmid ESX radiation is that it was driven by adaptation to accompanying gene content on the plasmid. To investigate this possibility, we analyzed gene content across related groups of plasmids (Table S4). Gene content on the plasmids was highly variable, and little to no gene content was uniquely shared among plasmids with similar ESX (plasmid gene content is discussed in more detail below). This indicates that divergence of plasmid-borne ESX is unlikely to have been driven by interactions between ESX and gene content mobilized on plasmids.

Bacteria can protect themselves from foreign DNA, including plasmids, using CRISPR-Cas nucleases. It is possible that plasmid ESX diverged in response to CRISPR found among mycobacterial host genomes. CRISPR-Cas systems have been identified previously in *M. tuberculosis, M. bovis,* and *M. avium* (He et al. 2012). We searched the annotations of 33 mycobacterial species for which finished genome sequence data were available, and only identified CRISPR loci in *M. canettii, M. kansasii, M. avium,* and the *M. tuberculosis* complex (MTBC). This indicates that plasmid-borne ESX divergence is unlikely to have been shaped by adaptation to host CRISPR, at least as they are currently recognized.

*Congruent evolution of plasmid conjugation machinery: T7SS and T4SS* As a final possibility, we investigated adaptation of plasmid conjugation systems as a driving force for divergence of plasmid ESX. Both T7SS and T4SS were found to be essential for plasmid conjugation in a recently discovered plasmid in *Mycobacterium marinum* (Ummels et al. 2014). With one interesting exception discussed below, we found T7SS to be invariably accompanied by T4SS in our plasmid sample, suggesting that their functions are interdependent across diverse mycobacterial plasmids.

Several plasmids found in *M. ulcerans* encoded an ESX 2P-like locus that was not invariably accompanied by a T4SS. Two other features distinguished these plasmids from those found in other species of mycobacteria. First, there were numerous transposable and other mobile elements on the plasmids, and second, the ESX locus showed evidence of progressive degradation with multiple, independent examples of loss of one or more genes within the locus (Figure 9).

Excluding *M*. ulcerans-associated plasmids, we found the core genes of mycobacterial ESX-encoding plasmids to consist of the T7SS genes (by definition), as well as T4SS genes *(virB4, tcpC)* and *espI,* which was in some cases located within the ESX locus and in others was located separately. These findings extend earlier observations of mycobacterial plasmids encoding an ESX-5 like locus (Ummels et al. 2014).

Individual phylogenies of T4SS and T7SS genes were congruent (Figure 8) and distinct from other gene content on the plasmids. This observation is consistent with the conjugation loci having a shared evolutionary history while other loci on the plasmid evolved independently. Congruence among T4SS and T7SS genes also suggests that the paralogous ESX systems trace to whole plasmid duplications. The combined T4SS/T7SS locus could in theory have been duplicated on individual plasmids, but the combined locus is ˜40kbp in size, and it seems unlikely that such a large duplication would be stable on a plasmid. In addition, we did not identify any plasmids with more than one ESX locus. A similar modular organization, with congruence among genes involved in conjugation, has been observed in other families of plasmids (Thomas 2000; Fernández-López et al. 2006).

### Adaptation of T7SS on the chromosome

Positive selection was evident on the ESX phylogeny in association with migration of the loci to the chromosome. This suggests that novel functions evolved for ESX following their incorporation into the chromosome. There is also evidence of positive selection along the branches separating various species of mycobacteria. This suggests that individual ESX systems may have functions that are specific to species or groups of species. Another possibility is that the advantageous mutations driving divergence of chromosomal ESX loci did not confer novel functions, but were advantageous as a result of interactions with loci elsewhere on the genome. Distinct functions have been identified for different ESX loci (i.e. ESX-1, -3, -5) and for the same loci in different species (e.g., ESX-1 in *M. tuberculosis* and *M. smegmatis),* indicating that at least in some cases the advantageous mutations conferred novel functions.

## Discussion

### Emergence of novelty on plasmids, complex dynamics among plasmids and chromosomes

Much of the prior research on gene duplication has focused on chromosomal duplications, either of the entire chromosome or one of its segments (Lynch & Conery 2000; Zhang 2003). The recent discovery of plasmid-borne ESX that are related to chromosomal systems (Ummels et al. 2014; Dumas et al. 2016; Newton-Foot et al. 2016) opens the possibility of a more complex evolutionary path underlying adaptation of the paralogous chromosomal loci. In addition to the previously described plasmid-borne lineages that root basal to ESX-1, -2, -3, and -5, we have identified a plasmid lineage that roots basal to ESX-4 and clarified the relationships among this ancestral group of loci (Figure 4). The finding that the earliest branching lineages on the ESX phylogeny are all plasmid-associated (Figures 3, 5B, 6) provides support for the hypothesis that the most recent common ancestor of these loci was plasmid-borne, and that divergence of the five major ESX lineages occurred on plasmids prior to their migration to the chromosome. Our proposed schematic outlining major steps during adaptation of the canonical ESX is shown in Figure 5A. The underlying history is necessarily very simplified in such a schematic, and the model is likely to be modified as further data become available.

The evolutionary history of mycobacterial ESX is evidently quite complex, with duplication and divergence occurring on plasmids, several migrations from plasmid to chromosome, lateral transfer among chromosomes (with or without a plasmid intermediary) as well as vertical inheritance, divergence on the chromosome and occasional loss of the loci from the chromosome. We saw evidence of ancient plasmid to chromosome migrations (e.g., of ESX-4 and -3 to the MRCA of mycobacteria; Figure 3) as well as more recent events (i.e. migration of ESX-N to *Nocardia* and ESX-4-bis to *M. goodii;* Figure 6). A similarly complex history has been observed previously, e.g. in IncW plasmids, where exchange of T4SS genes with the chromosome has occurred on several occasions (Fernández-López et al. 2006).

### Selective forces driving duplication and divergence of plasmid ESX

This complex history provides an interesting new paradigm for the evolution of novelty following gene duplication. Our analyses suggest that ESX duplication and divergence occurred on plasmids, and that this divergence was driven by positive selection. The initial event underlying creation of novel ESX loci appears to have been whole-plasmid duplication. Recent work in *Yersinia pestis* identified a positively selected phenotype associated with increased plasmid copy number (Wang et al. 2016). Positive selection for increased gene dosage may have similarly enabled the initial plasmid duplications underlying diverse T7SS. Such selection could operate at the level of the host cell, as in *Y. pestis,* or the plasmid, if, for example, it resulted in more efficient transfer of one or more plasmid copies.

We found that the T4SS and T7SS evolved in concert on the plasmids, along with *espI. EspI* has been shown to regulate ESX-1 in *M. tuberculosis* (Zhang et al. 2014); given its apparent essentiality in ESX encoding plasmids, we speculate it could play a similar role regulating plasmid-borne ESX. Diversification of these conjugative loci did not appear to have been driven by adaptation to different host species, CRISPR-Cas systems, or the gene content delivered by the plasmids. A possible alternative selection pressure is that imposed by plasmid incompatibility systems: i.e. the conjugation machinery differentiated to prevent conjugation between cells harboring incompatible plasmids. Surface exclusion is mechanistically distinct, but related to plasmid incompatibility and could also drive and maintain divergence of associated plasmid loci (Paulsson 2002; Garcillán-Barcia & de la Cruz 2008). Discriminatory transfer to host cells that lack incompatible plasmids is predicted to increase the fitness of the discriminatory plasmid (Paulsson 2002). Gene content on the plasmids was highly variable, suggesting that there is frequent recombination among them. Our finding that T7SS, T4SS and *espI* behave like a single locus (Figure 8), with little evidence of intra-locus recombination, provides further evidence that differentiation of these systems is maintained by selection, such as would be imposed by a plasmid incompatibility regime. Further studies are needed to investigate this hypothesis.

ESX-encoding plasmids in *M. ulcerans* provide an interesting example of apparent relaxation of selection to maintain conjugation machinery, with progressive degradation of the locus evident in extant plasmids (Figure 9). The *M. ulcerans* ESX plasmids also encoded the gene for mycolactone, which is essential for causing the ulcerative disease associated with *M. ulcerans* infection (George et al. 1999). Selection for plasmid-delivered gene content can stabilize non-transmissible plasmids (San Millan et al. 2014). We hypothesize that selection on *M. ulcerans* to maintain mycolactone-encoding plasmids relaxes selection on the plasmid to maintain its own conjugative machinery.

### Adaptation of ESX to the chromosome: initial stabilization by self-transfer?

We found evidence of directional selection - i.e. acquisition of specific advantageous mutations - in ESX following their migration to the chromosome (Figure 7). These advantageous mutations are the mechanism by which novel functions for T7SS would have been acquired. It is also possible that the new ESX duplicated the function of existing loci and that the mutations occurred because of epistasis, i.e. co-adaptation with other loci on the genome. Increased gene dosage is thought to be an important mechanism by which gene duplications are selected, and this could plausibly enable the retention of newly acquired loci that duplicated the functions of existing chromosomal loci (Bergthorsson et al. 2007; Bershtein & Tawfik 2008; Sandegren & Andersson 2009; Andersson & Hughes 2009). In the case of ESX, however, the loci diverged on the plasmids prior to their migration to the chromosome. While it is possible that further divergence of the migrant loci enabled functional convergence among chromosomal loci, this scenario seems quite complex and distinct functions have already been identified for some loci.

Both plasmid-borne and chromosomal ESX have been shown to mediate conjugation (Flint et al. 2004; Coros et al. 2008; Gray et al. 2013; Ummels et al. 2014; Gray et al. 2016). We found evidence suggesting that the chromosomal loci have been laterally transferred among bacterial species. Since ESX can mediate its own lateral transfer, it raises an interesting potential solution to Ohno's dilemma. Ohno's dilemma is the problem of how duplicate genes survive in the genome long enough to acquire mutations conferring a novel function, given the instability and potentially deleterious impacts of duplication (Bergthorsson et al. 2007). The situation here is slightly different in that the duplicate chromosomal loci arose *via* transfer from plasmids, but the question remains of how these migrant loci were stabilized on the chromosome long enough to co-adapt with it and evolve novel functionality.

We speculate that the migrant ESX loci acted initially as selfish genetic elements mediating their own transfer among chromosomes. We found *espI* and T4SS genes retained in association with more recent migrations, suggesting that all of the plasmid conjugation machinery was transferred initially, with subsequent remodeling of the locus. These duplicated, laterally spreading chromosomal ESX loci would provide a large genetic target for adaptive mutations conferring a new function (Bergthorsson et al. 2007). Fixation of these mutations would have been hastened by their lateral spread if the loci retained the capacity to mediate LGT. Whether they retained this ability or not, benefits provided by novel mutations would favor retention of ESX and resolve potential conflicts between the locus and its host genome.

### Functional diversification of chromosomal ESX

Our analyses of directional selection on chromosomal ESX delineate groups of loci that are likely to have differentiated from each other because of the acquisition of new functions. Functions have been identified for just a small number of ESX loci, and our results can aid further research in this area. For example, we found that *M. tuberculosis* ESX-3 is closely related to *M. marinum* ESX-3, without evidence of directional selection in the branches separating them (Figure 7, Figure S4). This suggests that *M. marinum* is likely to be a useful model for the study of ESX-3 functions in *M. tuberculosis*. The same is true of ESX-4, whereas *M. kansasii* may be a good model for *M. tuberculosis* ESX-2. ESX-1, which is an important virulence locus (Pym et al. 2002), appears to perform functions that are unique to *M. tuberculosis,* as does ESX-5. Experimental results from ESX-5 mutants in *M. tuberculosis* and *M. marinum* are consistent with our observations, since they suggest that this locus performs different functions in these closely related species (Shah & Briken 2016).

### Summary

The paralogous ESX loci are the product of a complex evolutionary history during which mycobacteria capitalized on diversity found among plasmid loci and repurposed the loci to perform diverse functions. This is an interesting paradigm for the generation of novelty via gene duplication, and such complex dynamics between mobile and core genomes may be important for other bacteria as well. Positive selection has played an important role in diversification of these loci, and we propose two potential solutions to the problem of how the duplicate loci were maintained long enough to acquire novel, adaptive mutations. Selection for increased plasmid gene dosage may have fostered the plasmid duplications, whereas we propose that an initial (or stable) LGT function may have favored retention of chromosomal loci following their migrations from plasmids. Delineation of this evolutionary history aids our understanding of the generation of evolutionary novelty and we propose ways in which these results can guide the choice of model organism and functional studies of these loci in *M. tuberculosis*.

## Acknowledgments

We thank Andrew Kitchen (University of Iowa) for his input on the manuscript. This material is based upon work supported by the National Science Foundation Graduate Research Fellowship Program [grant number DGE-1256259] and the National Institute of Health National Research Service Award [grant number T32 GM07215] to TDM. CSP is supported by National Institutes of Health [grant number R01AI113287].

